# An R-R-type MYB transcription factor promotes nonclimacteric pepper fruit ripening pigmentation

**DOI:** 10.1101/2022.09.15.507774

**Authors:** Ningzuo Yang, Jiali Song, Changming Chen, Binmei Sun, Shuanglin Zhang, Yutong Cai, Xiongjie Zheng, Bihao Cao, Guoju Chen, Dan Jin, Bosheng Li, Jianxin Bian, Jianjun Lei, Hang He, Zhangsheng Zhu

## Abstract

Carotenoids act as phytohormones and volatile compound precursors that influence plant development and confer characteristic colours, affecting both the aesthetic and nutritional value of fruits. Carotenoid pigmentation in ripening fruits is highly dependent on developmental trajectories. Transcription factors incorporate developmental and phytohormone signalling to regulate the biosynthesis process. In contrast to the well-established pathways regulating ripening-related carotenoid biosynthesis in climacteric fruit, carotenoid regulation in nonclimacteric fruit is poorly understood. Capsanthin is the primary carotenoid of nonclimacteric pepper (Capsicum) fruit; its biosynthesis is tightly associated with fruit ripening, and it confers red pigment to the ripening fruit. In this study, using a weighted gene coexpression network and expression analysis, we identified an R-R-type MYB transcription factor, DIVARICATA1, and demonstrated that it is tightly associated with the levels of carotenoid biosynthetic genes (CBGs) and capsanthin accumulation. DIVARICATA1 encodes a nucleus-localized protein that functions primarily as a transcriptional activator. Functional analyses demonstrated that DIVARICATA1 positively regulates CBG transcript levels and capsanthin contents by directly binding to and activating the CBG promoter transcription. Furthermore, the association analysis revealed a significant positive association between DIVARICATA1 transcription level and capsanthin content. Abscisic acid (ABA) promotes capsanthin biosynthesis in a DIVARICATA1-dependent manner. Comparative transcriptomic analysis of DIVARICATA1 in pepper and its orthologue in a climacteric fruit, tomato, suggests that its function might be subject to divergent evolution among the two species. This study illustrates the transcriptional regulation of capsanthin biosynthesis and offers a novel target for breeding peppers with high red colour intensity.

## Introduction

Carotenoids are a group of isoprenoid pigments that are synthesized by all photosynthetic organisms and some nonphotosynthetic fungi and prokaryotes (Hassan *et al*., 2019). Carotenoids are essential for plant photosynthesis, contributing to light harvesting and protecting plants from photooxidation (Karlova *et al*., 2014), and they are also precursors of certain phytohormones, namely, strigolactones (SLs) and abscisic acid (ABA) (Al-Babili and Bouwmeester, 2015). Carotenoid pigments are present in the fruits and flowers of many plant species, attracting animals for seed dispersal and pollination (Stanley and Yuan, 2019) and contributing to the aesthetic properties and perception of product quality. Humans cannot synthesize carotenoid pigments *de novo* but need them as important nutrients due to their potent antioxidant properties and/or provitamin A activity and (Zheng *et al*., 2020). Industrially, carotenoids are used across the cosmetic, food, health, feed, and pharmaceutical sectors (Bouvier et al., 2003, Noviendri et al., 2011). In plants, carotenoids synthesized in plastids from IPP are generated from the methylerythritol-4-phosphate (MEP) pathway (Sun et al., 2018). Two molecules of geranylgeranyl pyrophosphate (GGPP) are condensed by phytoene synthase (PSY), forming phytoene in the rate-limiting step of carotenoid biosynthesis (Berry *et al*., 2019). Subsequently, in the head, phytoene undergoes isomerization, epoxidation, and cyclization through phytoene desaturase (PDS), ζ-carotene desaturase (ZDS), carotenoid isomerase (CRTISO), lycopene β-cyclase (β-LCY), β-carotene hydroxylase (β-CH), zeaxanthin epoxidase (ZEP), and violaxanthin deepoxidase (ZDE) to antheraxanthin and violaxanthin (Berry *et al*., 2019; Rodriguez-Uribe *et al*., 2012). Ripening fleshy fruits have evolved to be attractive to frugivores to enhance seed dispersal and have become an indispensable part of the human diet (Karlova *et al*., 2014). The ripening developmental program of fleshy fruits involves changes in texture, flavor, aroma, and colour (Klee and Giovannoni, 2011). Carotenoids are regarded as among the most important pigments and are closely associated with the ripening pigmentation of many fruits. Many economically important fruits produce copious carotenoids during ripening and therefore experience the transcriptional control of carotenoid biosynthetic genes (CBGs) during fruit ripening. The fruit ripening process is generally classified as climacteric or nonclimacteric according to the physiological changes that occur (Li *et al*., 2022; Brumos, 2021). While ABA and ethylene are the primary phytohormones involved in the ripening of many fleshy fruits, their regulatory roles differ significantly between well-studied climacteric and nonclimacteric fruits (Li *et al*., 2022; Xiao *et al*., 2020). Ethylene plays a clear role in climacteric fruit ripening, whereas nonclimacteric fruit ripening is generally associated with ABA (Fenn and Giovannoni, 2021). The foremost model for carotenoid regulation is tomato, whose fruits are climacteric and in which ethylene biosynthesis and signalling are necessary for the onset and maintenance completion of ripening (Fenn and Giovannoni, 2021; Karlova *et al*., 2014). In tomato, the transcription factors NONRIPENING (NOR), COLORLESS NONRIPENING (CNR), RIPENING INHIBITOR (MADS-RIN), TOMATO AGAMOUS-LIKE1 (TAGL1), APETALA2a (AP2a) and FRUITFULL (FUL1 and FUL2), in concert with ethylene, regulate the ripening process, including modulating carotenoid accumulation (Klee and Giovannoni, 2011; Karlova *et al*., 2014). In addition, many recent studies have shown that ABA regulates climacteric fruit ripening and carotenoid accumulation by regulating transcription factors for ethylene biosynthesis and signalling, indicating that ABA-ethylene cross-talk occurs in climacteric fruit (Fenn and Giovannoni, 2021). ABA is generally acknowledged as one of the main regulatory phytohormones that acts during the ripening of nonclimacteric fruits (Fenn and Giovannoni, 2021). In recent studies, it has emerged that ABA modulates transcription-factor-controlled ripening-related processes in nonclimacteric fruit (Jia *et al*., 2016). This has been especially well studied in strawberries, in which ABA activates transcription factors that trigger a series of ripening processes, including anthocyanin accumulation, increased vacuolar hexose concentration, and decreased fruit acidity and chlorophyll content (Zhou *et al*., 2021). However, how ABA signalling modulates the activity of transcription factors that control carotenoid levels in nonclimacteric fruit is poorly understood.

Pepper (*Capsicum*) belongs to the Solanaceae family, and the genus *Capsicum* contains approximately 35 species, five of which have been domesticated: *C. annuum, C. baccatum, C. chinense, C. frutescens* and *C. pubescens* (Carrizo García *et al*., 2016). Capsicum fruit is typically considered nonclimacteric, but climacteric fruit ripening behaviour of some hot peppers has also been observed. Due to the desirable pigments, flavours, and aromas of its fruits, pepper is cultivated and distributed worldwide (Sun *et al*., 2022a; Zhu *et al*., 2019; Villa-Rivera and Ochoa-Alejo, 2021), including approximately 4.1 million tons of dry pepper and 54.9 million tons of green pepper (FAO). Mature pepper fruits express a diverse range of carotenoids, which confer different colours (Lee *et al*., 2021). Compared to other types of fruit, most ripe pepper fruits are red in colour. The diesters capsanthin and capsorubin confer red color to the fruits and account for 80% of the carotenoids in high-intensity accessions (Berry *et al*., 2019). The enzyme capsanthin-capsorubin synthase (CCS), a homologue of the lycopene cyclizing enzyme LCYb, catalyses the formation of capsanthin and capsorubin from antheraxanthin and violaxanthin in pepper fruit, representing species-specific carotenoid biosynthesis pathways in plants (Kim *et al*., 2014). In addition to providing colouration to pepper fruits, capsanthin is also widely used in the food additive, cosmetics, and pharmaceutical industries for its versatile properties (Hassan *et al*., 2019; Arimboor *et al*., 2015). Over decades of plant physiology and biochemistry research, including multiomics approaches, the *Capsicum* capsanthin biosynthetic pathways have been largely elucidated (Berry *et al*., 2019; Ou *et al*., 2018). With genetic mapping, homology-based cloning, and multiomics approaches, the genes that encode enzymes for carotenoid biosynthesis have been cloned and functionally identified from pepper (Popovsky and Paran, 2000). Many candidate genes whose mutants are associated with pepper ripening and fruit colour have been cloned and determined to encode biosynthetic enzymes. A mutation in the pepper *y* locus leads to a yellow colour of ripe fruit due to a loss of function of the *CCS* gene (Ha *et al*., 2007). The *c2* locus, mutations of which result in orange or peach colour in mature pepper fruit, cosegregates with *PSY*. Recently, a splicing mutation in *ZEP* leading to a premature stop codon was shown to contribute to orange colouration and altered carotenoid contents in pepper fruit (Lee *et al*., 2021). The capsanthin biosynthesis process is strictly controlled both spatially and temporally, and the CBGs are mainly transcribed in fruit from the break stage to the mature stage (Kim *et al*., 2014). The transcription levels of *PDS, PSY* and *CCS* are consistently higher in peppers with highly intense red colouration than in cultivars with low red intensity (Berry et al., 2019; Ha et al., 2007; Rodriguez-Uribe et al., 2012), indicating that capsanthin content is regulated at the transcriptional level. Upregulation of key CBGs in pepper seems to significantly increase the content of carotenoids, especially capsanthin (Rodriguez-Uribe *et al*., 2012; Berry *et al*., 2019). Moreover, CBG expression could be induced by the exogenous application of ABA, implying that ABA mediates certain transcription factors to regulate these CBGs (Xiao *et al*., 2020). However, the transcriptional regulatory mechanism of carotenoid biosynthesis in pepper is still unknown. Thus, the identification and functional characterization of the related transcription factors is imperative.

MYB transcription factors compose one of the largest families of transcription factors; these proteins share the conserved MYB DNA-binding domain (Zhu *et al*., 2019) and are categorized into subfamilies depending on the number of conserved MYB domain repeats. Plant MYB proteins are classified into three major groups: R2R3-MYB, with two adjacent repeats; R1R2R3-MYB, with three adjacent repeats; and a heterogeneous group collectively referred to as MYB-like proteins, which usually but not always contain a single MYB repeat (Dubos *et al*., 2010). In contrast, R2R3-MYB has been widely shown to play a role in controlling the biosynthesis of specialized metabolites such as lignins (Goicoechea *et al*., 2005), anthocyanins (Sun *et al*., 2016), glucosinolates (Sønderby *et al*., 2010) and capsaicinoids (Sun *et al*., 2022b; Zhu *et al*., 2019). R2R3-MYB transcription is emerging as a modulator of carotenoid production; for example, MlRCP1 in *Mimulus lewisii*, AdMYB7 in *Actinidia deliciosa*, MtWP1 in *Medicago* (Meng et al., 2019), and CaMYB306 in *Capsicum annuum* (Ma *et al*., 2022a) positively control carotenoid accumulation, whereas CrMYB68 performs the opposite function, repressing α- and β-branch carotenoids in *Citrus reticulata* (Zhu *et al*., 2017). Based on phylogenetic analysis, MYB-like proteins can be divided into five subgroups: CIRCADIAN CLOCK ASSOCIATED 1 (CCA1)-like, CAPRICE (CPC)-like, TELOMERIC DNA-BINDING PROTEIN (TBP)-like, I-box-binding-like and R-R-type (Gao *et al*., 2017). Compared to the well-studied R2R3 -MYB transcription factors in plants, only a few of these MYB-like genes have been functionally characterized (Arce-Rodríguez *et al*., 2021). Recent studies revealed that R-R-type MYB transcription factors integrate hormone biosynthesis and/or signalling to mediate responses to environmental challenges and cues, particularly in relation to ABA signalling (Fang *et al*., 2018; Guo *et al*., 2016). The R-R-Type MYB transcription factor AtMBS 1 (also known as AtMYBL) is involved in promoting leaf senescence and modulates an abiotic stress response in *Arabidopsis* (Zhang *et al*., 2011). AtDIV2, an R-R-type MYB transcription factor of Arabidopsis, negatively regulates salt stress by modulating ABA signalling (Fang *et al*., 2018). SRM1, a salt-related MYB1, can directly regulate the expression of NCED3/STO1, a key ABA biosynthesis gene, resulting in altered ABA levels in the overexpression lines (Wang *et al*., 2015). In rice, *MID1* (*MYB Important for Drought Response 1*) encodes an R-R-type MYB-like transcription factor that promotes male rice development under drought by modulating the expression of drought-related and anther developmental genes (Guo *et al*., 2016). Moreover, R-R-type MYB transcription factors have also been suggested to regulate development in other plant species. In *Antirrhinum majus*, the R-R-type MYB-like protein AmDIVIRICATA was proposed to regulate floral dorsoventral asymmetry (Gao *et al*., 2017; Perez-Rodriguez *et al*., 2005). Likewise, in *Plantago lanceolata*, the R-R-type MYB-like transcription factor PlDIV was reportedly co-opted to regulate cell proliferation during the early stages of pollen development (Reardon *et al*., 2014). This suggests that R-R-type MYB-like transcription factors execute diversified functions in different plant species. However, whether the R-R-type MYB-like transcription factor is mediated by ABA and the control of carotenoid biosynthesis in fleshy fruit remain unknown.

The phytohormone ABA is a combinatorial ripening transcription factor that controls pepper fruit capsanthin biosynthesis at the transcriptional level (Xiao *et al*., 2020). The availability of a large amount of RNA-seq data for pepper provides an ideal opportunity for investigating the regulation of capsanthin biosynthesis. In this study, through coexpression analysis, we identified and characterized the R-R-type MYB-like transcription factor DIVARICATA1 and validated its role in capsanthin biosynthesis. ABA could induce *DIVARICATA1* expression, and its expression pattern has undergone divergent evolution in pepper and tomato. This study identifies the first key regulator of capsanthin biosynthesis, establishing a foundation for further research investigating the regulatory network of capsanthin.

## Results

### An R-R-type MYB coexpressed with associated CBGs

Plant production of spatial, temporal or condition-specific metabolites and transcription factors serve as orchestrators of coordinated biosynthetic pathway genes (Sun *et al*., 2022; Sun *et al*., 2020; Sun *et al*., 2019; Zhu *et al*., 2019). This phenomenon can be leveraged by exploring RNA sequencing (RNA-seq) data to discover regulator genes that coexpress structural genes. Capsanthin levels appear to be influenced by the ontogenetic trajectory of the fruit. Capsanthin begins to accumulate at the fruit developmental stage of the mature green stage and peaks at the mature stage. Accordingly, the key genes *PDS, PSY* and *CCS* are also expressed in the fruit from the mature green stage to the mature stage. The pepper landrace CM334 (*C. annuum*) transcriptional data at seven different developmental stages were retrieved from previous studies (Kim *et al*., 2014). To identify potential transcription factors involved in the carotenoid pathway, especially capsanthin biosynthesis, we performed WGCNA. Among the identified modules, we identified the ModuleGID_darkgreen module as having an expression pattern tightly correlated with capsanthin biosynthesis (Fig. 1a). In total, 334 genes were included in this module, including *CCS (CA00g33800*) (kME *P* value=1.30E-09), which encodes the final enzyme in the pathway responsible for generating capsanthin and capsorubin. In addition, we found the gene *CA12g06700*, which encodes an R-R type MYB transcription factor belonging to the DIVARICATA family, referred to in this study as *DIVARICATA1*, which was tightly associated with the ModuleGID_darkgreen module (kME *P* value = 1.02E-08) (Fig. 1c and Supplementary Table 1). Moreover, *DIVARICATA1* was found to be highly coexpressed with CBGs. Pearson’s correlation coefficient analysis indicated that the *DIVARICATA1* transcription level had a strong positive correlation with *CCS* (r^2^ = 0.81), *PSY* (r^2^ = 0.81), *β-CH1* (r^2^ = 0.71), *PDS* (r^2^ = 0. 67), *PDS* (r^2^ = 0. 67), *ZDS* (r^2^ = 0.65) and *β-LCY1* (r^2^ = 0.59) (Fig. 1c-), respectively. We speculate that DIVARICATA1 might be involved in capsanthin biosynthesis by regulating certain carotenogenesis genes.

**Fig. 1.**
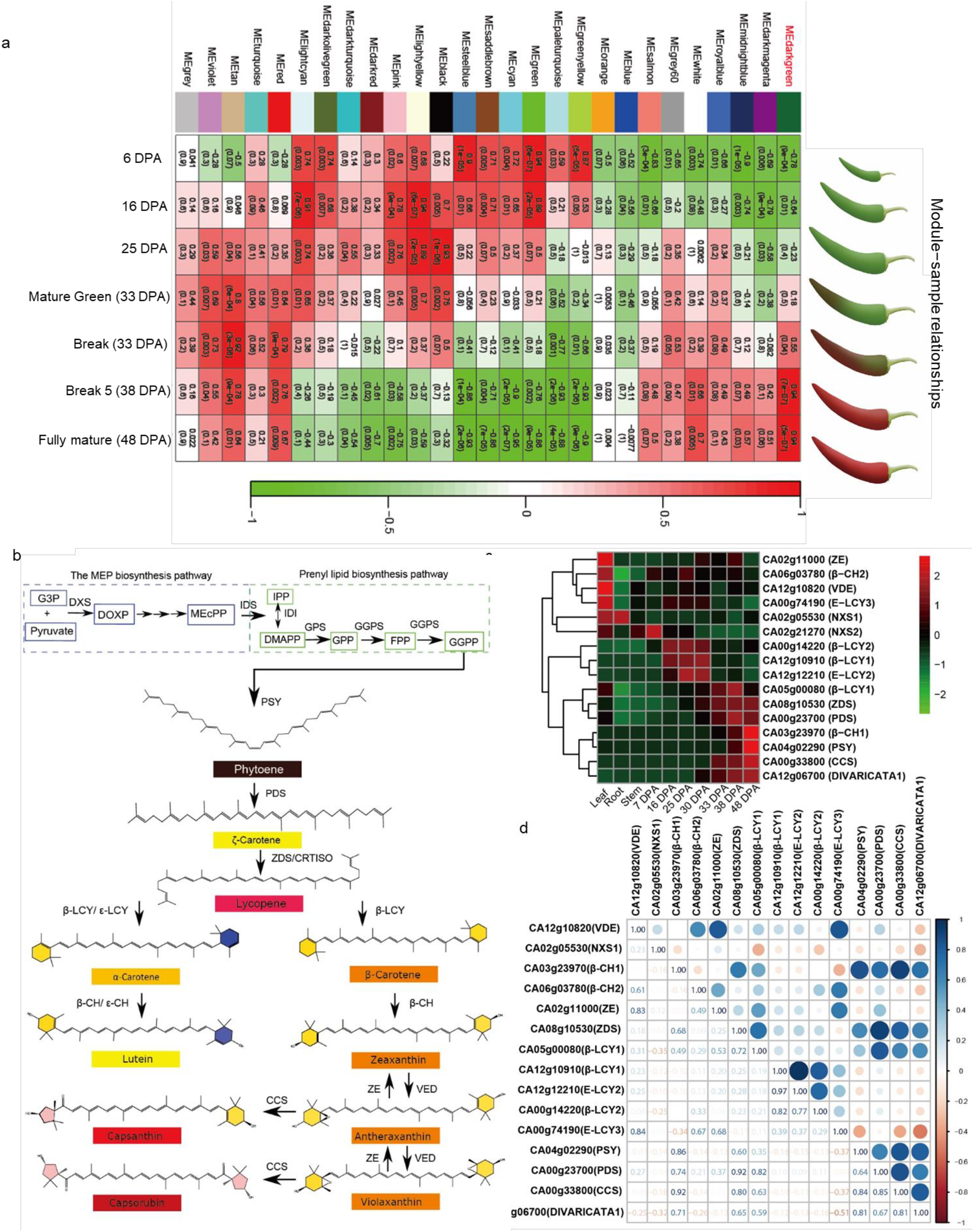
Carotenoid biosynthesis pathway and gene expression in pepper. (a) Module–fruit development stage association. Each row corresponds to a module. Each column corresponds to the cultivar CM334 at a certain developmental stage. The colour of each cell at the row–column intersection indicates the correlation coefficient between the module and the fruit developmental stage. A high degree of correlation between a specific module and developmental stage is indicated by red. The MEdarkgreen module, with the highest correlation between carotenoid biosynthesis and gene expression, is shown in red. (b) An illustration of the carotenoid biosynthesis pathway derived from previous studies, with minor modifications (Berry et al., 2019). (c) Heatmap showing CBG and DIVARICATA1 gene expression in pepper cultivar CM334. Days postanthesis (DPA) indicates the developmental stage of the fruit pericarp. (d) Pearson’s correlation coefficients for the gene transcription levels. The genes in different tissues and different developmental stages of the fruit pericarp were applied for analysis.

### DIVARICATA1 is associated with carotenoid biosynthesis in pepper fruit

To illustrate *DIVARICATA1* transcription and carotenoid biosynthesis, we determined the capsanthin content in the fruit pericarp of the elite ‘59’ inbred line (*C. annuum*) at six developmental stages (16, 25, 30, 33, 38, and 48 DPA) (Fig. 2a). The capsanthin in the 16 DPA immature fruit was at an undetectable level (Fig. 2b). Its abundance gradually increased during the mature green stage (25 to 30 DPA), with a rapid increase from 33 DPA to 38 DPA and a peak at 48 DPA. Real-time quantitative reverse transcription PCR (qRT–PCR) assays were performed to relate increased carotenogenesis in the ripening process to specific *DIVARICATA1* and carotenoid biosynthetic genes (Fig. 2c). The expression pattern of *DIVARICATA1* and CBGs was displayed in a tissue- and development-dependent manner. In the vegetative tissues, *DIVARICATA1* and CBGs were weakly expressed and almost undetectable. In contrast, *DIVARICATA1* was detected at a high transcription level in the pericarp from 30 DPA to 48 DPA, and the key carotenogenesis genes *PSY, β-CH1*, and *CCS* were also highly expressed at these stages. In the break stage, some genotypes displayed a uniform pattern of colouration (Fig. 2D). The results indicate that DIVARICATA1 expression in the fruit was consistently higher in the high red intensity part than in the low red intensity part (Fig. 2E). Taken together, these findings indicate that *DIVARICATA1* might regulate the expression of these genes during carotenogenesis.

**Fig. 2.**
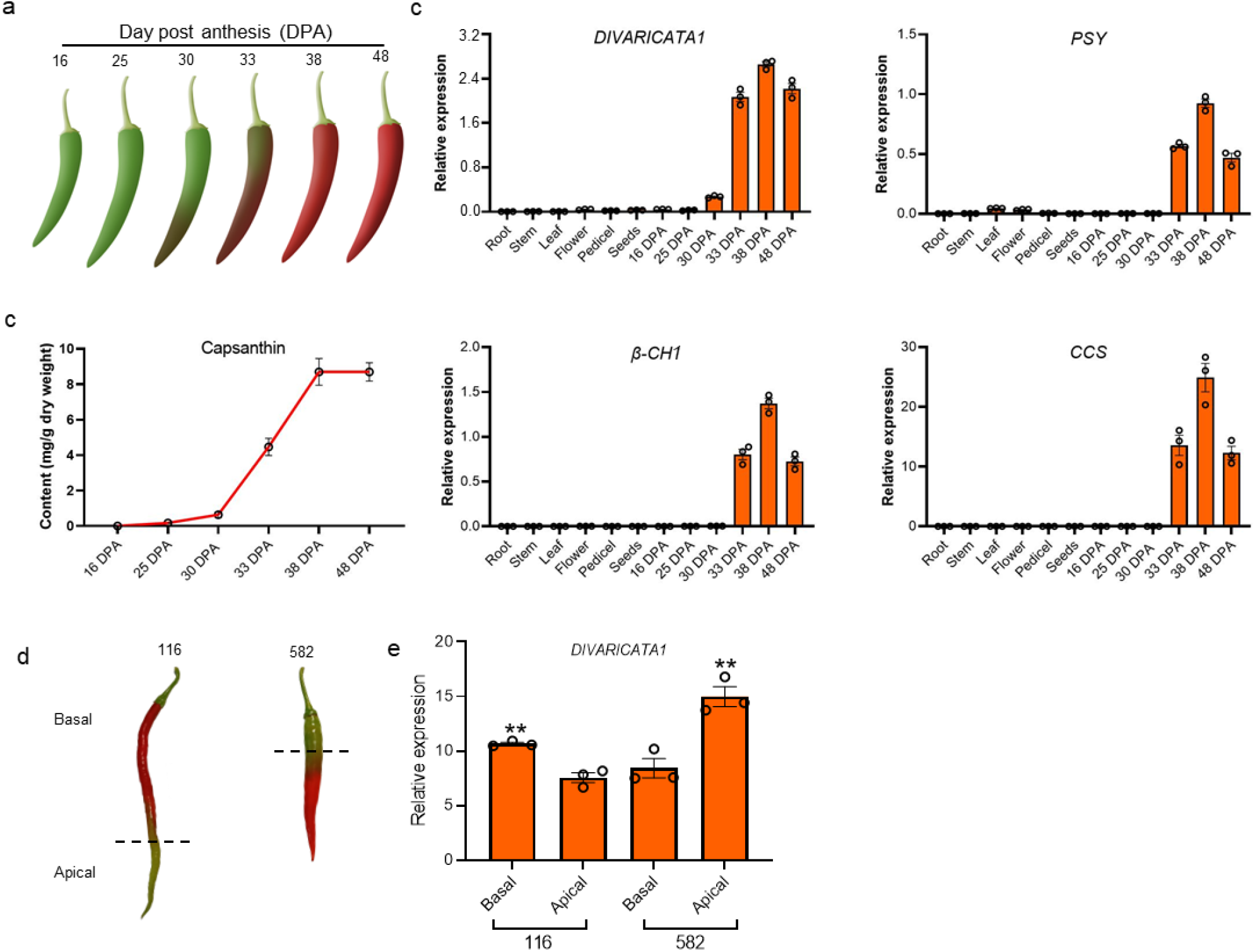
*DIVARICATA1* expression is associated with capsaicinoid biosynthesis. (a) Schematic diagram of pepper fruits at different developmental stages. (b) The contents of capsanthin in the fruit pericarp at different developmental stages. The fruit pericarp of the ‘59’ inbred line at different stages was used for analysis. Data are expressed as the mean± SE (*n* = 3). (c) Transcript levels of *DIVARICATA1* and CBGs in different tissues. Roots, leaves, stems, and apical buds were collected from 30-day-old seedlings. Fully blossomed flowers were collected from 60-day-old plants. Seeds and pedicel were collected from 38 DPA fruit, and the pericarp was collected from fruits at different developmental stages. The transcription of the target gene was expressed relative to the expression of the ubiquitin extension gene (*CA12g20490*). Data are expressed as the mean± SE (*n* = 3). (d) Fruits of *C. annuum* inbred line 116 and inbred line 582 at 33 DPA. (e) The transcription level of *DIVARICATA1* in apical and basal fruit of inbred line 116 and inbred line 582 as indicated in (D). Data are expressed as the mean ± SE (*n* = 3).

### DIVARICATA1 functions as a nucleus-localized transcription activator

The full-length coding sequence of *DIVARICATA1* was cloned from the ‘59’ inbred line. Nucleotide sequence analysis indicated that the coding sequence of *DIVARICATA1* was 669 bp in length, and the deduced amino acid sequence was 222 amino acids (Fig. 3a). To identify potentially conserved motifs in DIVARICATA1, the amino acid sequences from *Solanum lycopersicum, Arabidopsis thaliana* and *Antirrhinum majus* that shared the highest sequence identity with DIVARICATA1 were retrieved from public databases and aligned. As a result of the alignment, we found two DNA-binding domains (R domain) and the typical ‘SHAQKY/F’ motif, one of the main characteristics that distinguish the R2R3 MYB proteins of the DIVARICATA subfamily (Fig. 3a). To dissect the potential function of DIVARICATA in pepper, a phylogenetic analysis was carried out. The DIVARICATA-type MYB transcription factors from other plant species that shared the highest sequence identity with DIVARICATA were retrieved from public databases. In addition, some functionally identified MYB transcription factors that regulate carotenoids were also analysed. Within the DIVARICATA clade, the most closely related proteins from Solanaceae families was StDIVARICATA (*Solanum tuberosum*), SlDIVARICATA1 (*Solanum lycopersicum*), and NtDIVARICATA (*Nicotiana benthamina*), which shared a sequence identity of more than 85%. However, the functions of MYB factors in these Solanaceae species are unknown (Fig. 3b). Transcription analysis of the tomato cultivar ‘Heinz’ revealed that *SlDIVARICATA1* was rarely expressed in all tissues. The most closely related DIVARICATA transcription factor in *Arabidopsis thaliana* was AtDIVARICATA2, which showed 61% sequence identity and was previously found to negatively regulate salt stress by modulating ABA signalling (Fang *et al*., 2018). The *Arabidopsis thaliana* R-R-type MYB-like transcription factors AtMBS1 and AtMBS2 are involved in promoting leaf senescence and modulate an abiotic stress response (Zhang *et al*., 2011), sharing 46% and 51% identity with pepper DIVARICATA1, respectively. The protein in rice most closely related to DIVARICATA1 was MID1 (MYB Important for Drought Response 1), with 48% sequence identity. MID1 has been proposed to mediate ABA signalling to enhance the response to drought and other abiotic stresses (Guo *et al*., 2016). *Antirrhinum majus* AmDIVARICATA is a key factor involved in dorsoventral asymmetry (Galego and Almeida, 2002; Perez-Rodriguez *et al*., 2005), and it shares 53% sequence identity with DIVARICATA1. The R2R3 MYB transcription factors ElRCP1 (Sagawa *et al*., 2016), AdMYB7 (Ampomah-Dwamena *et al*., 2019) and RhPCP1 (Li *et al*., 2020) execute functions in regulating carotenoid biosynthesis; however, they shared only 33%, 41%, and 22% sequence identity with DIVARICATA1, respectively. These results indicate that the DIVARICATA function has undergone divergent evolution among different species and might have a novel function in *Capsicum* species to some extent.

**Fig. 3.**
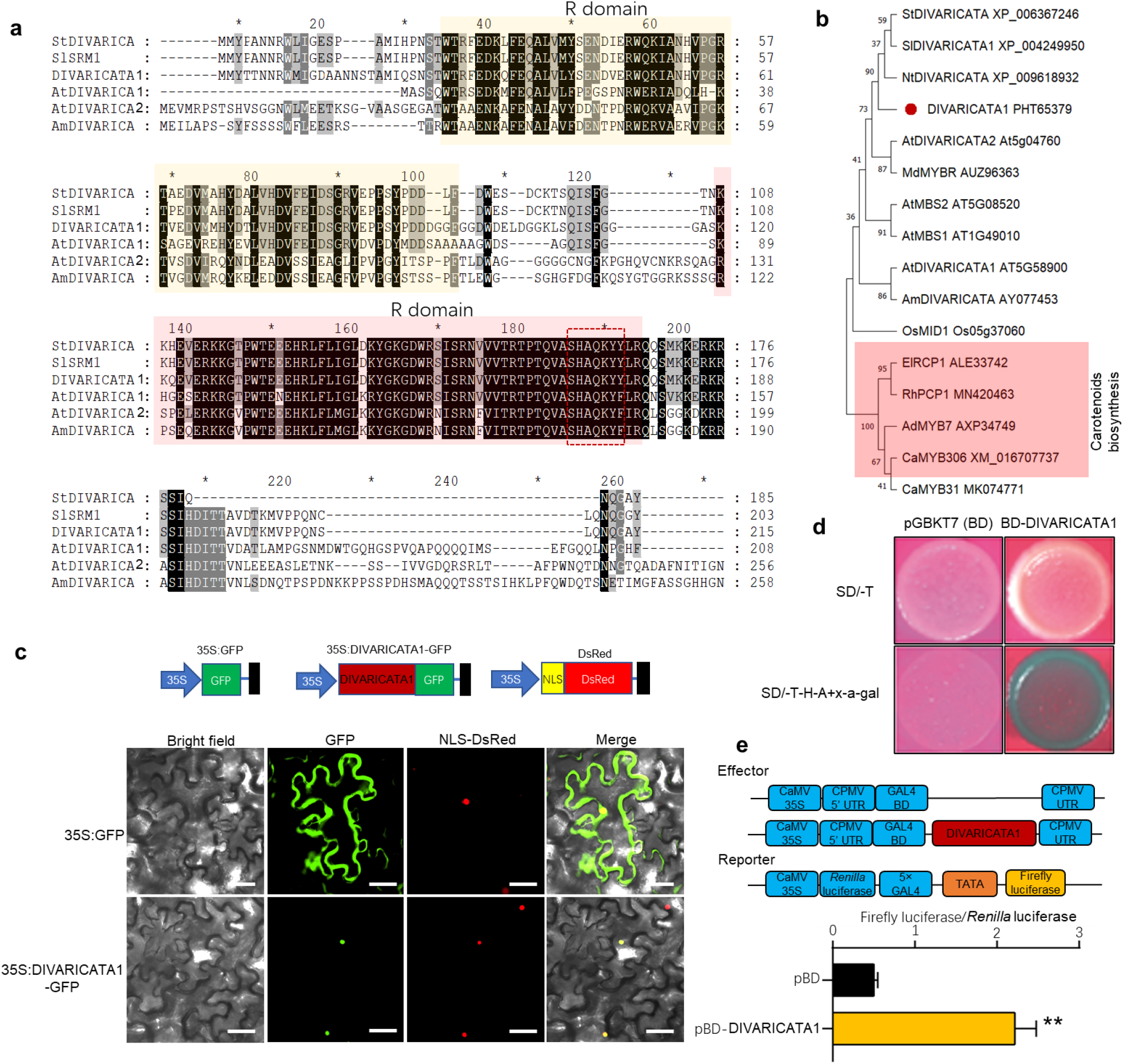
Sequence and transcriptional activity analysis of DIVARICATA1. (a) Sequence analysis of DIVARICATA and other MYB transcription factors. To compare the MYB sequences among different plant species, DIVARICATA1 was used as the search protein. MYB transcription factors sharing highly similar sequences in *Solanum lycopersicum, Solanum tuberosum, N. tabacum*, and *Arabidopsis thaliana* were retrieved from a public database. (b) A phylogenetic tree of MYB transcription factors. The evolutionary history was inferred using the neighbour-joining method. The percentage of replicate trees in which the associated taxa clustered together in the bootstrap test of 1000 replicates. The evolutionary distances were computed using the JTT matrix-based method and in units of the number of amino acid substitutions per site. Evolutionary analyses were conducted in MEGA11. (c) DIVARICATA1 is localized in the nucleus. *Agrobacterium tumefaciens* harbouring 35S:GFP or 35S:DIVARICATA1-GFP were injected into *N. benthamiana* leaves. NLS-DsRed served as a nuclear marker. Bar = 25 μm. (d) Activation analysis of DIVARICATA1 in yeast. Auxotroph plates (SD/–Leu–His–Ade–x-α-gal, below) showing activation of the protein. (e) Transcriptional activation process of DIVARICATA1 *in planta*. Data represent the mean± SD (*n* = 5). Student’s *t* test was used to identify significant differences compared to the empty vector control (\*\**P* < 0.01).

To investigate the subcellular localization of DIVARICATA1, the coding sequence of DIVARICATA1 was fused in frame with the GFP gene to generate a 35S:DIVARICATA1-GFP construct (Fig. 4c). The DsRed protein fused with a nuclear localization signal (NLS) served as the nuclear marker. Either 35S:GFP or 35S:DIVARICATA1-GFP was coexpressed with DsRed in *N. benthamiana* leaf epidermal cells. The DIVARICATA1-GFP fusion protein colocalized with DsRed (Fig. 4c), indicating that DIVARICATA1 functions as a nucleus-localized transcription regulator. To determine the transcriptional activation ability of DIVARICATA1, we performed transactivation activity assays *in vitro* and *in planta*. The DIVARICATA1 full-length coding sequence was fused in frame with the GAL4 DNA-binding domain, and the construct was transformed into the Y2H Gold yeast strain. We observed the growth of yeast carrying BD-DIVARICATA1 on SD/-Trp/-His/-Ade+x-α-gal medium, indicating strong activation activity (Fig. 3c). Next, we conducted a dual-luciferase (LUC) reporter assay to reveal whether this transcriptional activation ability is recapitulated in plants. GAL4 BD was fused in frame with the full-length coding sequence of DIVARICATA1 to generate the effector (Fig. 3e). The effector and reporter were coinfected into *N. benthamiana* leaves via *Agrobacterium-mediated* transformation. Compared with the control, DIVARICATA1 strongly enhanced firefly luciferase activity (Fig. 3e), indicating that DIVARICATA1 has transcriptional activity *in planta*.

**Fig. 4.**
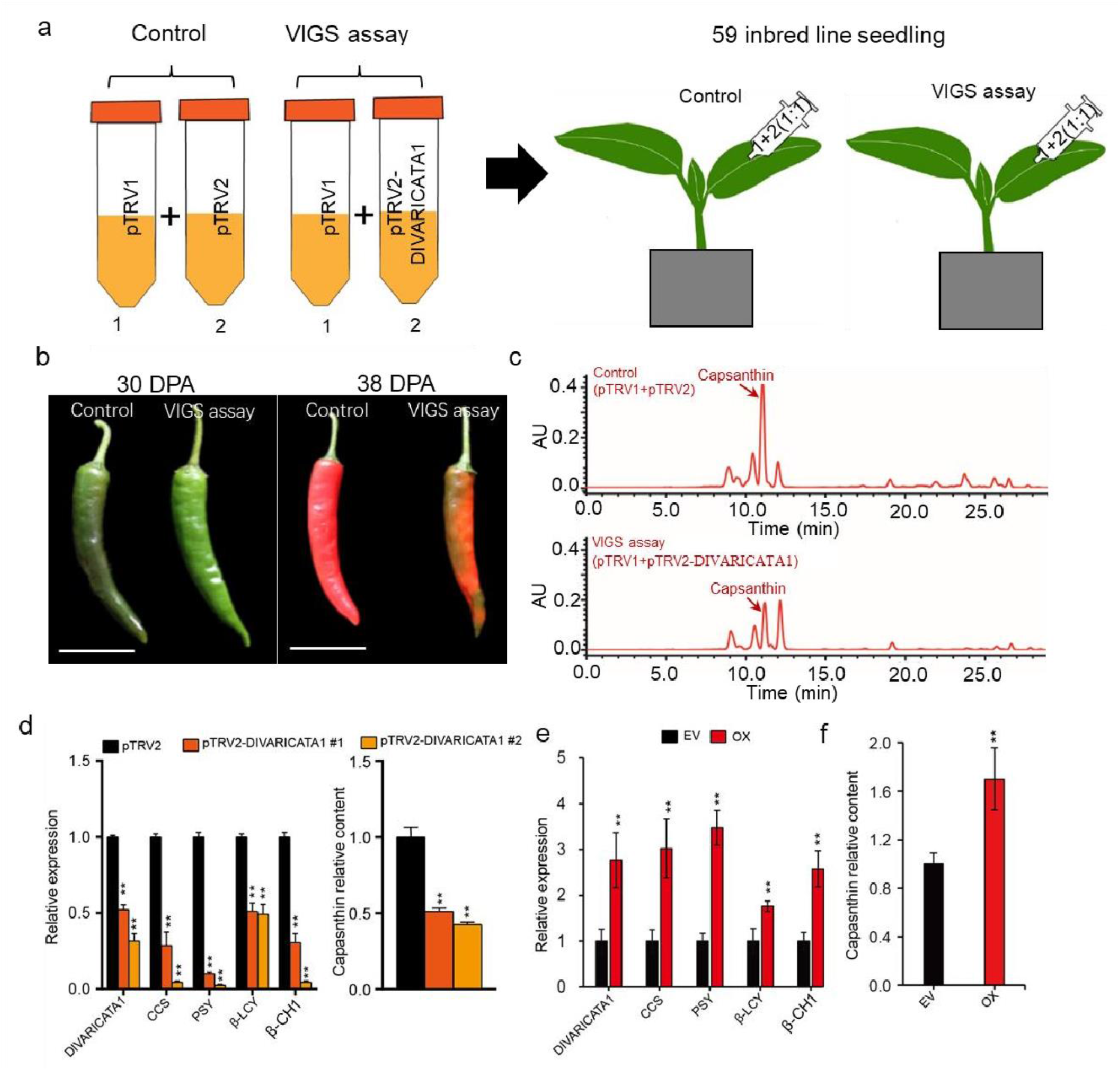
*DIVARICATA1* regulates CBG expression and capsanthin content. (a) Schematic diagram of the *DIVARICATA1* VIGS assay. *A. tumefaciens* harbouring empty vector (pTVR1+pTRV2) served as the control, and VIGS assay (pTVR1+pTRV2-*DIVARICATA1*) vectors were coinfiltrated (1:1) into 35-day-old seedling leaves of inbred line ‘59’. (b) The typical fruit phenotype at 30 DPA and 48 DPA. Bar = 5 cm. (c) Typical HPLC profile of carotenoids in control and VIGS assay ripe pepper fruit. The observation was performed at a wavelength of 470 nm. (d) Relative expression of *DIVARICATA1* and CBGs at 33 DPA in *DIVARICATA1* -silenced plants compared to control plants. The relative expression of the control was set to 1, and that of the *DIVARICA TA 1* -silenced plant was measured relative to that of the control. Data represent the mean ± SE (*n* = 3). Student’s *t* test was used to identify significant differences compared to the empty vector control (\*\**P*<0.01). (e) and (f) Transient overexpression of *DIVARICATA1* (OX) increases the transcription of CBGs and capsanthin content. The 30 DPA fruit was infiltrated with *A. tumefaciens* harbouring empty vector (EV) or *35S:DIVARICATA1* overexpression vector. The 38 DPA fruit pericarp was sampled for gene expression and capsanthin analysis. The relative CBG expression and capsanthin content of the EV were set to 1, and the expression in the OX plant was measured relative to that of the control. The data represent the mean± SD (*n* = 3). Student’s *t* test was used to identify significant differences compared to the EV control (\*\**P*<0.01).

### *DIVARICATA1* positively regulates CBG transcription levels and capsanthin contents

To demonstrate the function of DIVARICATA1 in peppers, we silenced *DIVARICATA1* in the inbred line ‘59’ via virus-induced gene silencing (VIGS) (Fig. 4a). Compared with the control silencing, *DIVARICATA1* significantly delayed the colouration time and decreased the red intensity (Fig. 4b). The decreased red intensity is consistent with the substantial decrease in capsanthin content observed in the mature stage (Fig. 4c). The silencing efficiency was determined by qRT–PCR assay of the *DIVARICATA1* transcript level in 38 DPA fruit pericarp. The *DIVARICATA1* transcription level in silenced lines was reduced to 50-35% of that of the control plants, which, consistent with CBG, was also significantly downregulated in silenced lines (Fig. 4d). Consequently, the capsanthin content in silenced fruit was decreased to 44-50% of that in the control (Fig. 4d; Supplementary Fig. 2). To confirm that DIVARICATA1 regulates capsanthin biosynthesis in pepper, we transiently overexpressed it in 30 DPA fruit and then analysed the 38 DPA fruit pericarp. We found that the DIVARICATA1 and CBG transcription levels increased 1.8–3.4-fold in the overexpressing (OX) fruits compared to the empty vector (EV) control (Fig. 4d), consistent with the 1.7-fold increase in capsanthin content in the overexpression fruit pericarp (Fig. 4f). In addition to the differences in carotenoids, we observed differences in the chlorophyll contents between silenced fruit and control fruit in our study (data not shown).

### DIVARICATA1 directly binds to and activates CBG promoter transcription

*DIVARICATA1* -coexpressed CBGs were regulated by DIVARICATA1, suggesting that these genes might be directly targeted by DIVARICATA1. Three key genes, *PSY*, *β-CH* and *CCS*, were selected for further analysis. We applied the DIVARICATA1 orthologous genes from *Arabidopsis thaliana* AtDIVARICATA1 as a Matrix ID to identify MYB-related binding *cis*-elements in the JASPAR database. We found that MYB-related binding *cis*-elements were present within the *PSY*, *β-CH1* and *CCS* promoter regions (Fig. 5a), suggesting that these genes could be targeted by DIVARICATA1. Luciferase reporter imaging showed that DIVARICATA1 enhanced the intensity of firefly luciferase signalling under the control of the *PSY*, *β-CH1* and *CCS* promoters (Fig. 5b), indicating that DIVARICATA1 could activate the transcription of these genes. Dual-luciferase reporter (DLR) system analysis of firefly LUC activity showed that the *PSY*, *β-CH1* and *CCS* promoters were strongly activated by DIVARICATA1 in *N. benthamiana* leaves (Fig. 5c). Next, we investigated how *CCS* was regulated by DIVARICATA1. In the Y1H system, we found that DIVARICATA1 associates with the *CCS* promoter in yeast (Fig. 5d). EMSA was adopted to confirm the binding of DIVARICATA1 to its MYB-related *cis*-element *in vitro*. The results showed that upon incubation with the DIVARICATA1 recombinant proteins, the migration of the bands of the wild-type probes was retarded, whereas upon incubation with mutated probes, no signal of binding was observed (Fig. 5c). To confirm the binding of DIVARICATA1 to the *CCS* promoter, we carried out ChIP–qPCR assays to compare the relative enrichment of the specific sequence of the *CCS* promoter. The DIVARICATA1 protein was associated with the *CCS* promoter in a region-dependent manner, and region 2, which contains two MYB-related binding *cis*-elements, had a higher enrichment level than region 1, indicating that DIVARICATA1 binds the *CCS* promoter (Fig. 5e). Together, these results suggest that DIVARICATA1 regulates capsanthin biosynthesis by directly regulating CBG expression.

**Fig. 5.**
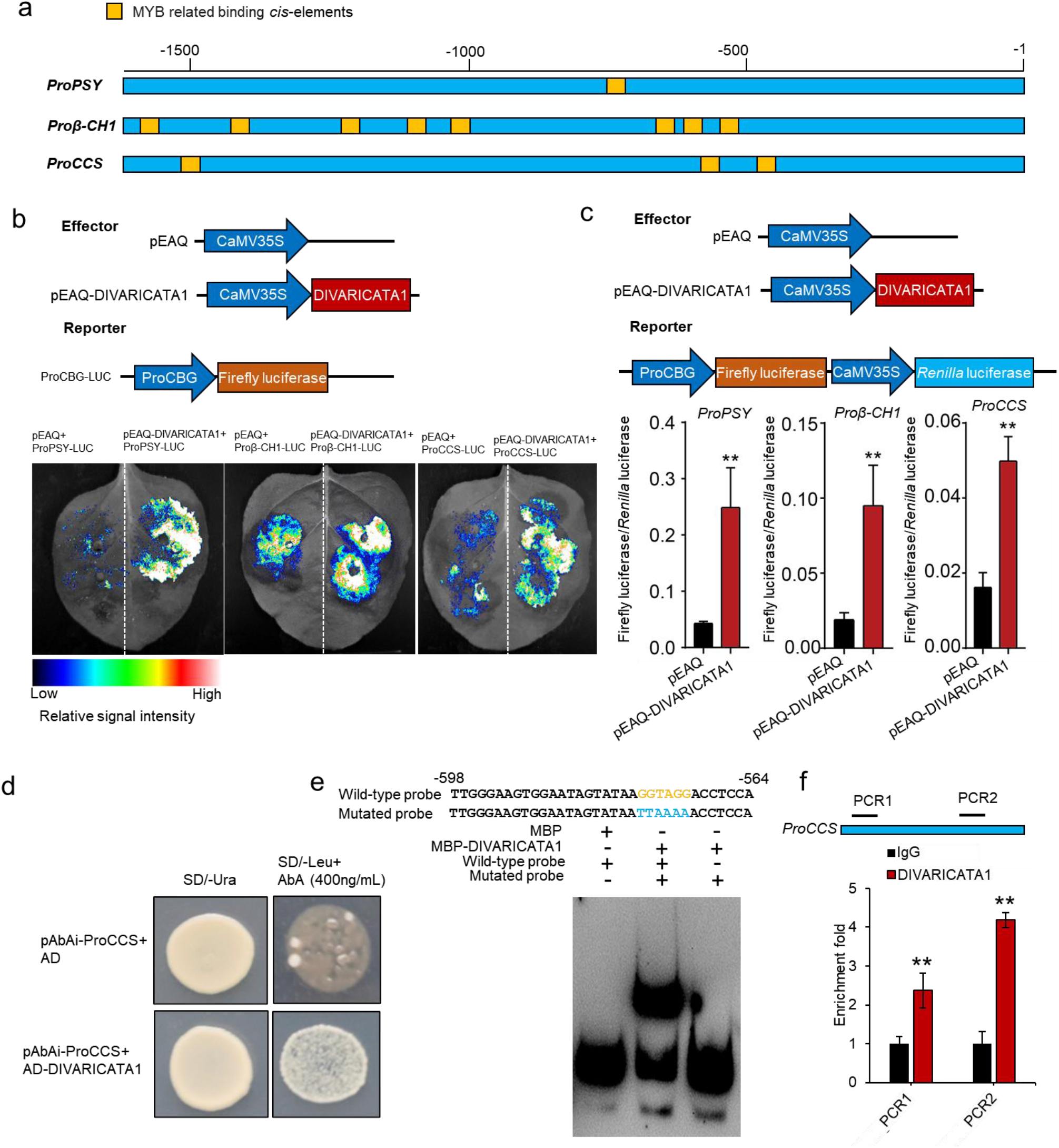
DIVARICATA1 targets the promoters of CBGs. (a) Schematic diagram of MYB-related binding *cis*-elements within the CBG promoter. The 1600 bp sequences upstream of the start codons of three CBGs (*PSY, β-CH1* and *CCS*) were analysed for MYB-related binding *cis*-elements via the JASPAR database. (b) DIVARICATA1 enabled the activation of *PSY, β-CH1* and *CCS* gene promoter transcription in *N. benthamiana* leaves. (c) DIVARICATA1 enabled the activation of *PSY, β-CH1* and *CCS* gene promoter transcription in *N. benthamiana* leaves. Data represent the mean± SD (*n* = 5). (d) DIVARICATA1 is associated with the promoter of the *CCS* gene in yeast. SD/-Ura medium lacking uracil; SD/–Leu medium lacking leucine; AbA, aureobasidin A. (e) EMSA of *in vitro* binding of DIVARICATA1 to the CCS promoter. (f) ChIP–qPCR analysis of DIVARICATA1 target to the CCS promoter. DNA was extracted from the pericarp of 38 DPA fruits and used for ChIP-PCR enrichment analysis. Values represent the mean ± SD (*n* = 3). Student’s *t* test was used to identify significant differences compared to the IgG control (**, *P* < 0.01).

### *DIVARICATA1* transcription level positively correlated with capsanthin content

The content of capsanthin in ripening *Capsicum* fruit determines the red intensity of the fruit (Berry *et al*., 2019). We selected twenty pepper accessions with different red intensities. High-performance liquid chromatography (HPLC) analysis of 38 DPA fruit carotenoids showed that capsanthin was distributed at variable levels among the twenty pepper accessions. The transcript profiles of *DIVARICATA1* and capsanthin in fruits of different accessions were measured, and linear regression analysis was performed. As shown in Fig. 6, the changes in capsanthin content correlated positively with those in *DIVARICATA1* transcripts (R = 0.75, *p* = 0.00012), suggesting that the variation in *DIVARICATA1* transcript levels is consistent with the observed variation in capsanthin content among different accessions.

**Fig. 6.**
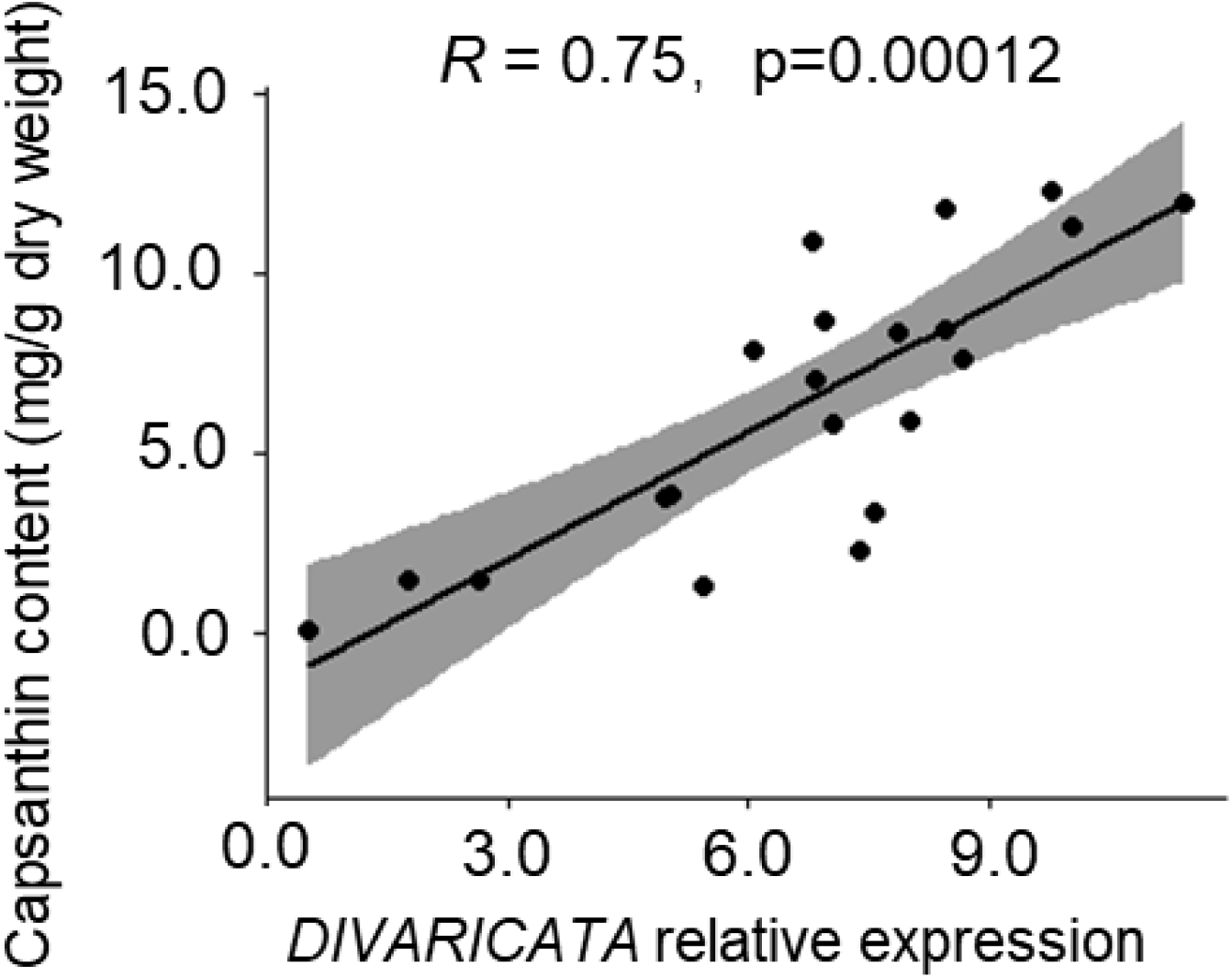
Correlation of *DIVARICATA1* expression and capsanthin content. Linear regression analysis between *DIVARICATA* expression and capsanthin content. The transcription of the target gene is expressed relative to that of the ubiquitin extension gene (*CA12g20490*). The capsanthin content of twenty pepper accessions was determined at 38 DPA. Significant differences were determined by R statistics.

### ABA-induced capsanthin biosynthesis is dependent on DIVARICATA1

Fleshy fruits are physiologically classified into climacteric and nonclimacteric groups. Pepper is a nonclimacteric fruit whose ripening is thought to be dependent on ABA (Tian *et al*., 2016; Xiao *et al*., 2020). In pepper fruits, an increase in ABA is associated with ripening, and carotenoid accumulation is thought to mainly be mainly dependent on ABA (Xiao *et al*., 2020). Considering that DIVARICATA1 plays an essential role in regulating carotenoid biosynthesis in pepper, we speculate that ABA signalling might act upstream of DIVARICATA1 in the regulation of capsanthin production. To confirm this hypothesis, mature green pepper fruits were treated with ABA and adopted for analysis. The results indicate that ABA induced the expression of *DIVARICATA1* and CBGs (i.e., *PDS, PSY, β-LCY1, β-CH1, CCS*) in a gene-dependent manner (Fig. 7 a-e and Supplementary Fig. 3). The highest expression levels of these genes were mainly observed from 3 h to 6 h after ABA stimulation, and the level of upregulation ranged from 2-fold to 5-fold compared to the initial level. Accordingly, the capsanthin level of ABA-treated 38 DPA pepper fruit was 40% higher than that of the control (Fig. 7 f), indicating that ABA could promote capsanthin production. Next, we investigated ABA-mediated DIVARICATA1 regulation of capsanthin production. The *DIVARICATA1* -silenced and control plant mature green fruits were treated with ABA. The results indicated that *DIVARICATA1* and CBGs were significantly induced by ABA in pTRV2 control pepper fruits (1.6-fold to 5.2-fold compared with those without ABA treatment), whereas these genes were only slightly induced by ABA in *DIVARICATA1*-silenced pepper fruits (1.3-fold to 2.0-fold compared with those without ABA treatment) (Fig. 7g). The gene expression was inconsistent with that in *DIVARICATA1* -silenced pepper, and the capsanthin content of ABA-treated fruit was only 15% higher than that of the control without ABA treatment (Fig. 7h). These results indicate that the ABA-mediated increase in capsanthin levels is largely dependent on *DIVARICATA1*.

**Fig. 7.**
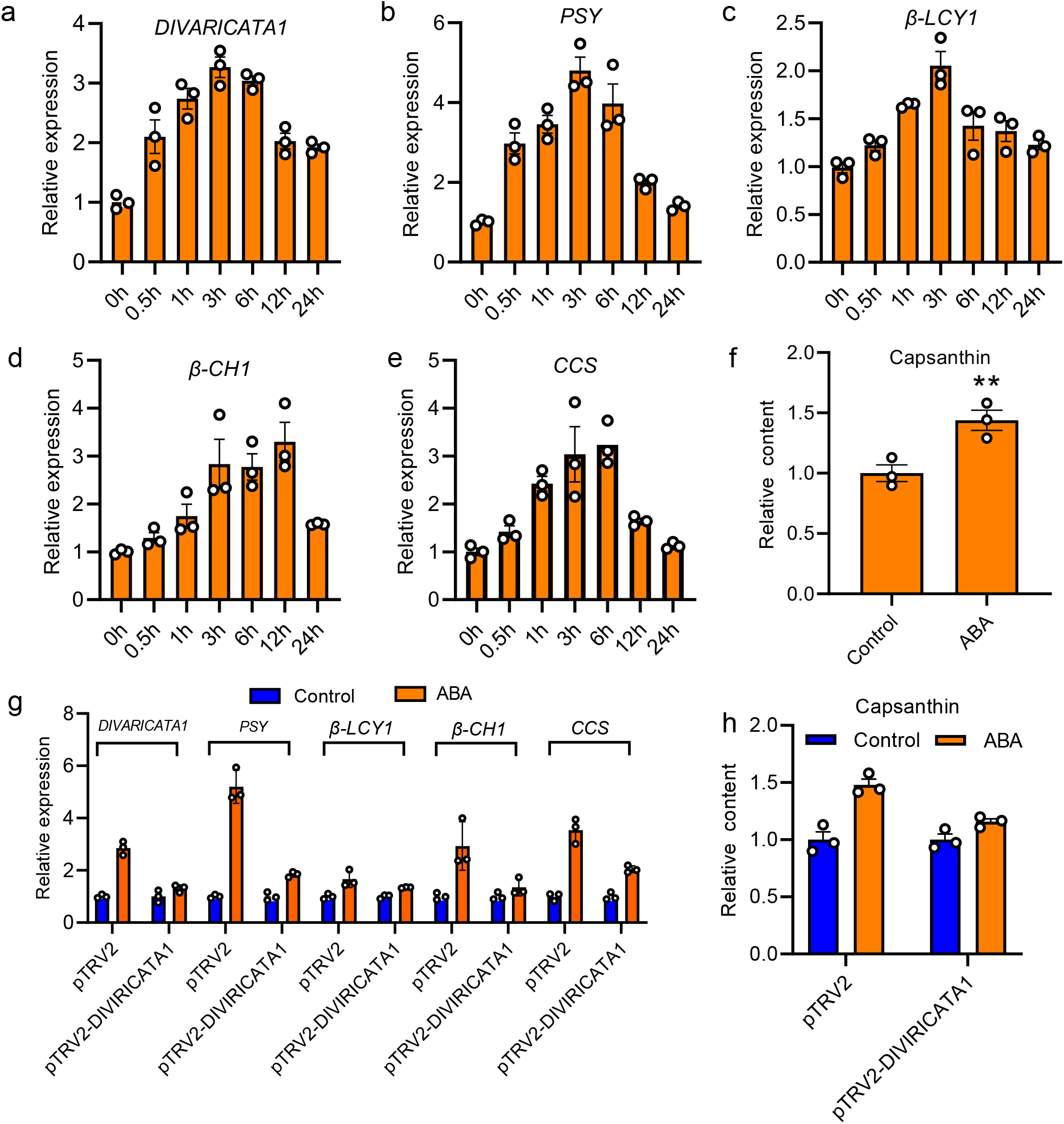
ABA-mediated capsanthin biosynthesis is dependent on *DIVARICATA1*. (a-e) Transcript levels of *DIVARICATA1* and CBGs after ABA treatment. Mature green pepper fruits were treated with 50 μmol ABA and sampled at the indicated time points. The initial target expression level at 0 h was set as 1, and expression levels at other time points were normalized to that at the initial stage. The data are expressed as the mean± SE (*n* = 3). (f) The capsanthin content of control and ABA-treated fruit. Data are expressed as the mean±SE (*n* = 3). Student’s *t* test was used to identify significant differences compared to the empty vector control (\*\**P* < 0.01). (g) Relative gene expression in control and *DIVARICATA1*-silenced fruit with or without ABA treatment. The expression of target genes corresponding to the control. Data represent the mean± SE (*n* = 3). (h) The capsanthin content of control and *DIVARlCATA1*-silenced fruit with or without ABA treatment. Data are expressed as the mean± SE (*n* = 3). Student’s *t* test was used to identify significant differences compared to the empty vector control (*\*\*P* < 0.01).

### Divergent evolution of DIVARICATA1 function in pepper and tomato

Pepper and tomato fruits are typical representatives of Solanaceae plants with nonclimacteric fruit and climacteric fruit, respectively. Despite their differences in ethylene biosynthesis and ethylene perception, both ripening processes are accompanied by the accumulation of carotenoids (Dong *et al*., 2014). In the climacteric fruit of tomato, ethylene plays an important role in triggering the onset of the ripening process, including the accumulation of carotenoids, particularly lycopene. However, in nonclimacteric pepper fruits, ripening is thought to be dependent on ABA (Supplementary Fig. 4 and Supplementary Fig. 5). Since DIVARICATA1 is essential in regulating carotenoid biosynthesis in pepper, we performed comparative transcriptomics to determine whether its orthologous function is conserved or divergent in tomatoes. We investigated orthologous genes previously identified in tomato ripening pigmentation to identify conserved and differentiated regulatory mechanisms between hot pepper and tomato. The expression of upstream genes, including *PSY*, *ZDS*, and *PDS*, was conserved during fruit ripening (Fig. 8a). In contrast, the downstream genes *β-CH1* and *CCS* showed distinct expression patterns in hot pepper and tomato. The *β-CH1* and *CCS* expression levels were extremely high during pepper ripening. *DIVARICATA1* was expressed at very low levels (FPKM lower than 30) during tomato ripening, whereas it was expressed at extremely high levels (FPKM higher than 1000) during pepper ripening, suggesting that DIVARICATA1 function evolved divergently in the two species. The conservation and divergence of this signalling pathway and transcription of these genes may lead to distinct outcomes (Fig. 8b).

**Fig. 8.**
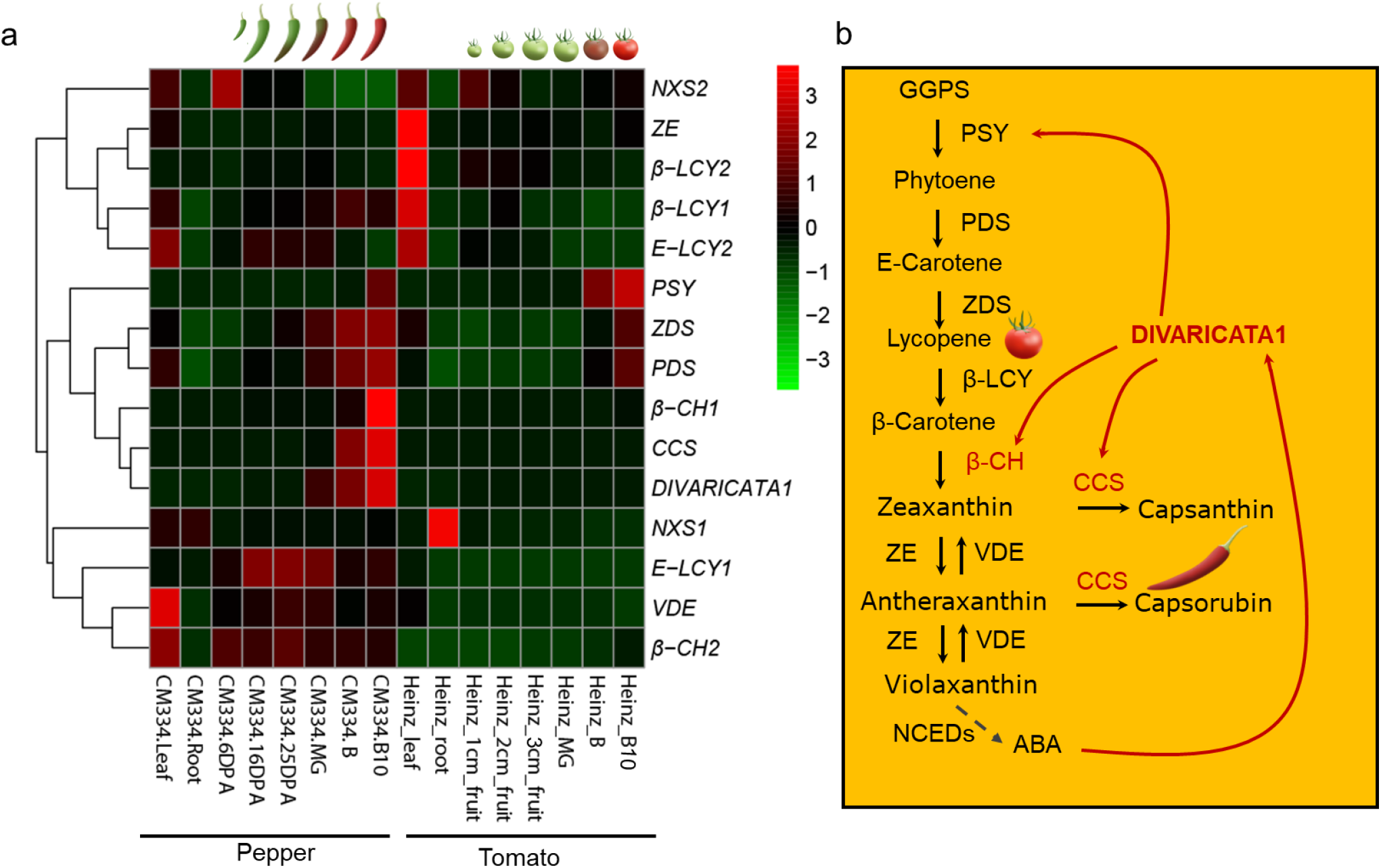
Transcriptional divergence and conservation of carotenoid-related genes in hot pepper and tomato. (a) Heatmap of carotenoid biosynthesis pathway genes and *DIVARICATA1*. The normalized RNA-seq data were retrieved from public databases. (b) Carotenoid biosynthesis pathway. The key enzymes marked with red indicate genes that are highly expressed in pepper but rarely expressed in tomato. The red line indicates genes that can be regulated by DIVARICATA1 in pepper.

## Discussion

### Pepper capsanthin biosynthesis is regulated at the transcriptional level

The biosynthesis of specialized metabolites in plants is often limited to specific tissues and coordinated developmental trajectories (Colinas and Goossens, 2018). During the ripening process, pepper fruit undergoes profound morphological and metabolic transformations as the chloroplast differentiates into a chromoplast due to the accumulation of ripening-related carotenoids (Berry *et al*., 2019). In fleshy fruits such as tomatoes and watermelon, high levels of the carotenoid lycopene accumulate, and early steps in the lycopene synthesis pathway are upregulated, but late cyclization is restricted (Fraser *et al*., 2002). In the fruits of red pepper, the content of the major chloroplast pigment lutein decreases during the ripening process from its highest level during the nonripe stage, whereas the levels of *β*-carotene gradually increase, and other *β*-carotenoids, including *β*-cryptoxanthin, zeaxanthin, and capsanthin, begin to be synthesized *de novo* (Rodriguez-Uribe *et al*., 2012; Song *et al*., 2020). In this study, in pepper fruit, accumulation of the major ripening pigment capsanthin was found to be associated with fruit development. The capsanthin biosynthesis and accumulation started at 25 DPA, and the peak content was found at 38 DPA (Fig. 3b). The capsanthin accumulation patterns observed in this study are consistent with previous reports that the highest levels of capsanthin could be detected at 7-10 days after breaking in pepper accessions with different red intensities (Berry *et al*., 2019). However, some other studies reported that pepper fruit started breaking colour at 45 DPA (Konishi et al., 2019). These different conclusions might be caused by differences in the genotypes and/or fruit development criterion scales among different studies. Consistent with capsanthin biosynthesis, key CBG expression was also found to be highly developmentally regulated in this study. In particular, *PSY*, *PDS* and *ZDS* encode enzymes for early steps in carotenoid synthesis, and *β-CH1* and *CCS* encodes enzymes for carotenoid cyclization, which ultimately results in the production of capsanthin, which is highly expressed from 30 DPA (breaking stage) to 48 DAP (ripening stage) (Fig. 1b and Fig. 2b). The expression levels of the *PSY, PDS, β-CH1* and *CCS* genes are high in peppers with high total carotenoid levels, whereas one or two of these genes are not expressed in peppers with lower total carotenoid levels (Ha *et al*., 2007; Rodriguez-Uribe *et al*., 2012; Kilcrease *et al*., 2015). The transcriptional regulation of carotenoid accumulation in pepper fruit continues to be a complex process in which the transcription of *PSY*, *PDS*, *β-CH1* and *CCS* vary between different red pepper varieties, and the contents of specific carotenoids varies as a result (Kilcrease *et al*., 2015). Nevertheless, *PSY* and *CCS* transcript levels were significantly positively correlated with capsanthin content in the ripening stage in accessions with different red intensities (Berry *et al*., 2019; Rodriguez-Uribe *et al*., 2012), indicating that the expression levels of the carotenoid biosynthetic genes are critical regulators of the high total carotenoid and capsanthin levels.

### DIVARICATA1 positively regulates fruit capsanthin accumulation during the ripening process

Compared to the understanding of the molecular basis underlying fruit ripening in climacteric fruits, including tomatoes (Brumos, 2021; Karlova *et al*., 2014), our knowledge of the regulation of ripening in nonclimacteric fruits such as pepper is still very limited. Pepper is a nonmodel but important horticultural crop, and the discovery of the nutritional value of capsanthin biosynthetic transcription regulators during the ripening process has been hindered. The regulation of specialized metabolite biosynthesis seems to be controlled at the transcriptional level, which is generally dependent on the interaction of DNA-related mechanisms and the activity of transcription factors that may act in a combinatorial manner (Colinas and Goossens, 2018). These master regulators appear to function as orchestrators in the coordinated control of key metabolic genes since they display similar expression patterns (Sun *et al*., 2022; Zhu *et al*., 2019). In this study, we adopted coexpression analysis to discover capsanthin biosynthesis regulators (Fig. 1). However, in the coexpression analysis of key genes, CBGs were unexpectedly grouped into different modules (i.e., *PSY* in ModuleGID_saddlebrown; *PDS* in ModuleGID_darkred; *ZDS* in ModuleGID_red; *β-CH1* in ModuleGID_cyan; *CCS* gene in ModuleGID_darkgreen). We speculate that the transcriptional regulation of carotenoid accumulation in pepper fruit is likely a complex process in which the expression of downstream factors is tightly associated with abscisic acid (ABA) biosynthesis. Most likely, a large set of transcription factors individually or synergistically target the CBG promoter to regulate carotenoid biosynthesis in pepper. This would be analogous to the case in tomato, in which a set of transcription factors from different families, such as RIN, TAGL, FUL1, FUL2, NOR, CNR and HB-1, individually or in different combinations bind different target genes to provide individual or overlapping contributions to the expression of CBGs (Stanley and Yuan, 2019). Compared to other specialized metabolites, in which the biosynthetic pathway genes and master regulator expression display a highly coexpressed pattern (Bai et al., 2022; Lau and Sattely, 2015; Nett et al., 2020), the expression of CBGs and their regulators may be more variable. Indeed, this could be learned from the observation that transcription factors, CBGs and regulators, were rarely identified through a coexpression analysis strategy. Among the reported results, a complete carotenoid pathway governed by one regulator is still rare. Nevertheless, in the ModuleGID_darkgreen module containing *CCS*, we identified a novel gene, *DIVARICATA1*, that encodes an R-R-type MYB transcription factor of unknown function (Supplement Table 1). Correlation coefficient analysis revealed that *DIVARICATA1* had an especially high positive correlation with key late CBGs (i.e., *CCS* and *β-CH1*) (Fig. 1 and Fig. 2), and we further demonstrated that it functions as a transcriptional activator by directly targeting CBG promoters to enhance capsanthin production (Fig. 3, Fig. 4 and Fig. 5). Moreover, the *DIVARICATA1* transcription level was significantly positively correlated with capsanthin content; presumably, *DIVARICATA1* is the capsanthin master regulator. However, we cannot rule out the possibility that other transcription factors in ModuleGID_darkgreen are associated with ripening pigmentation; for example, the tomato RIN orthologue *CaMADS-RIN (CA00g39870*) is also present in this module (Supplement Table 1). Overexpression of pepper *CaMADS-RIN* in *rin* mutant tomato plants can rescue the carotenoid accumulation defect phenotype, promote ethylene-dependent and ethylene-independent gene expression, and indicate that CaMADS-RIN affects tomato fruit ripening in both ethylene-dependent and ethylene-independent aspects (Dong *et al*., 2014), illustrating that MADS*-*RIN function is largely conserved in climacteric and nonclimacteric fruits and can be interchanged. It was reported that the pepper R2R3 MYB transcription factor CaMYB306 executes dual functions for accelerating fruit colouration and negatively regulates cold resistance (Ma *et al*., 2022). Unlike CaMYB306, which regulates upstream CBGs to regulate the total carotenoid contents in pepper and tomato, DIVARICATA1 seems to have evolved to preferentially regulate downstream CBGs (i.e., *CCS* and *β-CH1*) and govern capsanthin content. However, the underlying genetic mechanism causing the differential expression of DIVARICATA1 in accessions of different genetic backgrounds needs further study.

Interestingly, unlike the case for other specialized plant metabolites, such as flavonoids, anthocyanins and glucosinolates, the orthologous gene functions and regulatory mechanisms governing carotenoid synthesis are highly conserved across plants. Although a large set of transcriptional regulators of carotenoid biosynthesis in plants have been identified, functional consensus regulators are still limited. In kiwifruit (*Actinidia deliciosa*), a promoter screen identified the R2R3-MYB transcription factor AdMYB7 as a putative regulator of AdLCYB (Ampomah-Dwamena *et al*., 2019), and stable overexpression of this gene in tobacco increased the expression of the *NbPSY, NbPDS, NbZDS, NbLCYB, NbLCYE*, and chlorophyll biosynthesis genes, resulting in increased carotenoid content. The *Medicago truncatula* gene *MtWP1* encodes an anthocyanin-related R2R3-MYB activator that plays a critical role in regulating floral carotenoid pigmentation by directly regulating the expression of carotenoid biosynthetic genes including MtLYCe and MtLYCb (Meng *et al*., 2019). In the monkeyflower species *Mimulus lewisii*, an R2R3-MYB gene called *Reduced Carotenoid Pigmentation 1 (RCP1*) positively regulates all of the CBP genes expressed in flowers, contributing to the bright yellow colouration of the floral nectar guides (Sagawa *et al*., 2016). The *Citrus* transcription factors CsMADS5 and CsMADS6 positively modulate carotenoid metabolism by directly regulating carotenogenic genes (Lu *et al*., 2018; Lu *et al*., 2021). Papaya (*Carica papaya*) CpNAC1 was shown to bind directly to the NAC-binding sites of the *CpPDS2/4* promoters (Fu et al., 2016). In watermelon, a nonclimacteric fruit, the expression of tomato homologues SlCNR, SlAP2a, and SlERF6 was correlated with ripening and carotenoid biosynthesis (primary lycopene); however, that of MADS-RIN, TAGL1 and NAC-NOR was not. In this study, we demonstrated that the R-R-type MYB transcription factor DIVARICATA1 regulates carotenoid levels in peppers. This raises thought-provoking questions regarding why so many candidate regulators but so few consensus regulators exist. Perhaps it is not particularly surprising that carotenoids serve very different functions in different organs and that species seem to have evolved their own carotenoid regulators. Indeed, most carotenoid regulators were identified in ripening fruits, especially climacteric tomatoes (e.g., RIN, TAGL1, FUL1/FUL2, HB-1), as components of the ethylene signalling network (Karlova *et al*., 2014; Stanley and Yuan, 2019). Therefore, it would not be unexpected if these tomato fruit ripening genes do not regulate carotenoid biosynthesis in other tissue types or in nonclimacteric fruits. Tomato, the foremost model fruit for research, accumulates lycopene, whereas fruits of many other plant species accumulate abundant downstream products (Stanley and Yuan, 2019). Some carotenoid regulators, such as CpNAC1, CrMYB68, CsMADS5, CsMADS6, CsERF061, and AdMYB7, were verified to be associated with CBG gene promoters mainly through *in vitro* gel shift assays and/or DLR assays and further characterized by transient or stable overexpression in a heterologous host (Lu *et al*., 2018; Zhu *et al*., 2021; Lu *et al*., 2021; Li *et al*., 2017). Therefore, we should bear in mind that these regulators should be regarded as “candidates” instead of true carotenoid regulators until knockout or knockdown data become available.

### Divergent evolution of DIVARICATA1 function in plants

It seems that DIVARICATA function has undergone divergent evolution in plants. DIVARICATA1, a homologue of Arabidopsis AtDIVARICATA2, is universally expressed in tissues and is dramatically induced by exposure to salt stress and exogenous ABA; functional analysis showed that it negatively regulates salt stress by modulating ABA signalling (Fang *et al*., 2018). Regarding its orthologous protein in *Antirrhinum majus*, AmDIVARICATA, which is mainly expressed in petals, is a key factor by controlling the specification of particular cells within the ventral petal that adapt the corolla to specialized functions in pollination (Galego and Almeida, 2002). Likewise, expression and *cis*-regulatory element analysis found that the functions of AmDIVARICATA orthologs in orchids has been evolutionarily conserved (Valoroso *et al*., 2019; Valoroso *et al*., 2017). It has also been learned that an *Arabidopsis* loss-of-function mutant in *div2* led to elevated levels of ABA-related genes; *div2* was therefore proposed to be a transcription repressor (Fang *et al*., 2018). Among the DIVARICATA subgroup, the secondary close orthologues *AtMBS1 (AtMYBL*) and *AtMBS2* encode the R-R type MYB transcription factors, whose expression is significantly induced by ABA and salt stress (Zhang *et al*., 2011). Constitutive expression of the *AtMBS1* led to the upregulation of *Senescence-related gene (SRG*) and downregulation of *Ribulose-1,5-bisphosphate carboxylase small chain 1A (RBCS1*) and *Light-harvesting chlorophyll a/b-binding protein (LHB1B2*), which had significant influences on senescence parameters, including Chl content and membrane ion leakage. In parallel, the rice *MID1 (MYB Important for Drought Response 1*) was primarily expressed in root and leaf vascular tissues, with low levels in the tapetum, and was induced by drought and other abiotic stresses (Guo *et al*., 2016). *MID1*-overexpressing plants were more tolerant to drought at both the vegetative and reproductive stages and produced more grains under water stress but exhibited less severe anther defects. MID1 could bind to the promoters of two drought-related genes (*Hsp17* and *CYP707A5*) and one anther developmental gene (*KAR*). In tomato, the I-box binding factor LeMYBI protein contains a SHAQKYF amino acid signature motif in the second MYB-like repeat. LeMYBI binds specifically to the I-box sequence of the photosynthetic carbon dioxide fixation key enzyme *Ribulose-1,5-bisphosphate carboxylase small chain (RBCS1, RBCS2* and *RBCS3A*) gene promoters to activate its expression (Rose *et al*., 1999). It appears that changes in the *DIVARICATA* subgroup expression pattern resulted in its partly or completely divergent functional evolution among plant species. Although DIVARICATA executes diversified functions, the response to the ABA signal seems to be conserved to a large extent.

Although the sequence identity of DIVARICATA1 and its orthologues is highly conserved in Solanaceae plants, the functions of these orthologues in other Solanaceae plants have not been determined. Intriguingly, in this study, DIVARICATA1 was found to be highly expressed in pepper fruit from the break stage to the ripening stage, whereas its orthologue in tomato underwent divergent evolution. This outcome is perhaps not particularly surprising in that tomatoes are climacteric fruit, and thus, ethylene biosynthesis and signalling are necessary for the onset and completion of ripening, including the promotion of the accumulation of lycopene in mature fruits. During tomato fruit development, there are increases in the levels of the key ripening regulators MADS-RIN, FUL1, FUL2, TAGL1, CNR, NOR and AP2a, some of which serve dual functions for activation of ethylene biosynthesis and signalling, and the early CBGs (i.e., *SlPSY1* and *SlPDS*), whereas repression of *SlLCYE* and *SlLCYB* converts lycopene to other downstream products. In contrast, as pepper is a nonclimacteric fruit, ABA biosynthesis and signalling are crucial for the completion of ripening, including accelerating the accumulation primarily of the carotenoid capsanthin in ripening fruits (Fig. 7). During pepper and tomato fruit ripening, the expression of MADS-RIN, TAGL and NOR, which regulate upstream CBGs, is conserved (Kim *et al*., 2014), ensuring metabolic flux to lycopene. However, in pepper, perhaps genetic factors incorporating the ABA signal triggers species specific expression of DIVARICATA1, which activates the expression of the downstream cyclization enzyme genes *CCS* and *β-CH1* to ensure that the precursors are converted to capsanthin. However, it should be kept in mind that some other nonclimacteric fruits, such as watermelon, also accumulate lycopene, and orange accumulates abundant downstream products (e.g., β-carotene and xanthophylls). Whether DIVARICATA1 orthologues execute conserved or divergent functions in these species needs to be addressed. Apparently, the nonclimacteric fruit plant-specific evolution of *DIVARICATA1* mediates ABA signalling targeting downstream CBGs to govern metabolic flux conversion downstream, especially for capsanthin, at least in pepper (Fig. 8).

In this study, through coexpression analysis, we identified a gene module, ModuleGID_darkgreen, related to capsanthin biosynthesis. We demonstrated that DIVARICATA1 functions as a transcriptional activator that mediates ABA signalling to regulate capsanthin biosynthesis. The *DIVARICATA1* transcription level is tightly associated with fruit capsanthin content, and this information could facilitate the selection of high-content capsanthin pepper lines that harbour elite *DIVARICATA1* via haplotype base marker-assisted selection. The identification of *DIVARICATA1* provides a foundation for further research investigating the signalling and transcriptional regulatory networks for capsanthin biosynthesis in peppers. Furthermore, the discovery of DIVARICATA1 offers a new starting point for insights into the phytohormone onset of the transcriptional regulatory evolution of carotenoids in climacteric and nonclimacteric fleshy fruits.

### Experimental procedures

#### Plant materials

The elite pepper inbred line ‘59’ has high yield, resistance to diverse pathogens and good abiotic stress tolerance. To determine capsanthin content and analyse target gene expression in twenty different pepper accessions (inbred line ‘41’, ‘43’, ‘48’, ‘59’, ‘CA1’, ‘CA2’, ‘CA3’, ‘CA4’, ‘CA5’, ‘CA6’, ‘Cf1’, ‘Cf2’, ‘Cf6’, ‘678’, ‘732’, ‘LXJ1’, ‘YDH’, ‘BJ’, ‘HL13’ and ‘HL23’), seeds were sown in the nursery site, and 35-day-old seedlings were transplanted into 35-cm nonwoven pots. The plants were grown in a greenhouse with a daily temperature of 25-27 °C, night-time temperature of 20-22 °C, relative humidity of 60%, 16/8-h light/dark cycle, and light intensity of 6500 lux. After pepper fruit development to the selected stages, the fruits were sampled. The samples were frozen in liquid nitrogen and stored in a -80 °C freezer.

#### Determination of carotenoid contents

Frozen pericarp tissues were freeze-dried for 24 h with a freeze dryer (Labconco/Freezone, Labconco, America) and ground into fine powder by an A 11 basic analytical mill (IKA, Germany). The extraction of pepper carotenoids was performed as described previously (Song *et al*., 2020) with some modifications. Briefly, a total of 0.5 g of the freeze-dried samples was added to 8 ml of extracting solution containing hexyl hydride, acetone and absolute ethyl alcohol (2:1:1, HPLC grade), and then the samples underwent ultrasonic-assisted extraction for 30 min. Five millilitres of the supernatants were mixed with 5 ml of extraction solution and transferred to centrifuge tubes. Then, the mixtures were mixed with an equal proportion of saturated NaCl solution. The supernatants were transferred, and 2 ml of KOH and methyl alcohol (1:9) were added, mixed and then incubated for saponification for 12 h at room temperature. Finally, the extract was mixed with 2 ml of MTBE (methyl tert-butyl ether) and saturated NaCl solution. The supernatant was rinsed three times with saturated NaCl solution.

The analysis methods were described in a previous study (Zheng *et al*., 2019), with minor modifications. Mobile phase A (methanol/H2O, 9:1 [v/v]) and mobile phase B (100% [MTBE] containing 0.01% [w/v] butylated hydroxytoluene) were used for HPLC (Alliance E2695, Waters, America) at a flow rate of 1 ml/min. The gradient elution conditions were as follows: 0-6 min, 8–44% B; 6–11 min, 48–50% B; 11–14 min, 48–50% B; 14–30 min, 50-64% B; 30–34 min, 64-8% B; 34-36 min, 8% B. Detection was performed at 450 and 470 nm for capsanthin.

#### Sequence alignment and phylogenetic tree construction

The R-R-type MYB transcription factors showing the highest sequence identity with DIVARICATA1 and other well-characterized MYB transcription factors were retrieved from the NCBI (https://www.ncbi.nlm.nih.gov/) and Sol Genomics Network (http://solgenomics.net/) public databases and used for phylogenetic tree construction. The sequences were aligned by CLUSTALX 2 with the default parameters, and the phylogenetic tree was constructed by MEGA-X using the neighbourjoining method with 1000 bootstrap replications.

#### Quantitative real-time RT–PCR analysis

Total RNA was extracted from the samples using the HiPure HP Plant RNA Mini Kit (Magen, China). RNA was transcribed with the HiScript III 1st Strand cDNA Synthesis Kit (+gDNA wiper) (Vazyme Biotech, China). Quantitative real-time RT-PCR analysis was performed on a CFX384 Touch detection system (Bio-Rad, USA). As an internal control, the *ubiquitin extension gene (CA12g20490*) served as the reference gene (Liu *et al*., 2021). Each value represents the mean of three biological replicates. The primers used in this study are listed in Supplementary Table 2.

#### Identification of coexpression modules and visualization of gene expression

CM334 (*C. annuum*) is a Mexican landrace that displays red colour on ripening, and its genome sequence and transcriptome data were released in 2014 (Kim *et al*., 2014). These data include transcription data for genes from different tissues and fruits at different developmental stages. A gene coexpression network was built using the WGCNA package in R. The heatmaps display the expression of the levels of the assigned genes.

#### Subcellular localization

The full-length coding sequence of DIVARICATA1 was cloned into the pEAQ-GFP vector and fused with GFP. *Agrobacterium tumefaciens* GV3101 containing the corresponding constructs was infiltrated into 28-day-old *N. benthamiana* leaves. After *N. benthamiana* leaves were inoculated with *A. tumefaciens* for two days, the GFP signal was detected by confocal fluorescence microscopy (Carl Zeiss, Germany). NLS-DsRed served as the nuclear marker (Sun *et al*., 2020). Transcriptional activation analysis

A yeast transcriptional activation assay was performed as described previously (Zhu *et al*., 2019). The full-length CDS of *DIVARICATA1* was ligated into pGBKT7 to generate BD-DIVARICATA1. BD-DIVARICATA1 or the empty BD vector was then transformed into the yeast Y2H Gold strain and incubated on SD/-Trp medium at 30 °C for 3 days. The positive clones were picked and diluted in 0.9% NaCl solution, and 10 μL of each dilution was inoculated onto SD/-Trp-His-Ade medium. After 3-5 days, the clones were stained with x-α-Gal (Clontech, USA).

The transcriptional activation analysis of *DIVARICATA1 in planta* was carried out according to a previous study (Zhu *et al*., 2019; Sun *et al*., 2020). The full-length CDS of *DIVARICATA1* was cloned into the pEAQ-BD vector and fused with the GAL4 DNA-binding domain. Briefly, the *Agrobacterium tumefaciens* strain GV3101 containing the effector or pEAQ-BD and reporter was infiltrated into 28-day-old young *N. benthamiana* leaves. Three days after infection, the firefly luciferase and *Renilla* luciferase activities in the dual-luciferase reporter assay system (Promega, USA) were measured on a Varioskan™ LUX multimode microplate reader (Thermo Scientific, USA).

#### Y1H assay

The full-length coding sequence *DIVARICATA1* was cloned into pGADT7 to serve as the Y1H prey (AD-DIVARICATA1). The *CCS* promoter fragment containing a MYB-related *cis*-element was ligated into the pAbAi vector. The Y1H experiment was carried out according to the manufacturer’s protocol for the Matchmaker Gold Y1H system (Clontech, USA).

#### VIGS analysis

The coding sequence fragment of *DIVARICATA1* was amplified from the inbred line ‘59’. The fragment was cloned into pTRV2 to generate the silencing vector pTRV2-DIVARICATA1. VIGS was carried out according to previous studies with minor modifications (Zhu *et al*., 2019). Briefly, *A. tumefaciens* strain GV3101 harbouring the pTRV2-DIVARICATA1 and pTVR1 vectors was coinfiltrated into 35-day-old seedling leaves of the inbred line ‘59’. The empty vectors pTRV2 and pTVR1 served as controls. Total RNA was isolated from 38 DPA fruit pericarp for gene expression analysis and capsanthin analysis.

#### Dual-luciferase reporter assay

The upstream start codons of *CCS* (1843 bp), *PSY* (1917 bp) and *β-CH1* were PCR amplified from inbred line ‘59’ genomic DNA and cloned into pGreenII-0800 to generate reporters. The full-length CDS of *DIVARICATA1* was cloned into the pEAQ vector to generate an effector. *Agrobacterium tumefaciens* GV3101 containing the pEAQ-DIVARICATA1 effector in combination with the corresponding reporters was infiltrated into 28-day-old *N. benthamiana* leaves. After incubation for three days, firefly luciferase and *Renilla* luciferase were measured in a dual-luciferase reporter assay (Promega, USA). The relative activity was expressed as the ratio of firefly luciferase to *Renilla* luciferase.

#### Transient Expression Analysis

Transient overexpression analysis was carried out according to previous studies with minor modifications (Sun *et al*., 2019). Briefly, *A. tumefaciens* GV3101 harbouring the empty vector pEAQ or pEAQ-DIVARICATA1 at an optical density of 0.6 was infiltrated into 30 DPA fruits of the inbred line ‘59’ via the peduncle with a syringe. Each fruit was injected with approximately 0.6 mL of *A. tumefaciens* to avoid fruit drop. After 8 days of infection, the pericarp was sampled for RNA extraction for gene expression analysis and capsanthin measurements.

#### Electrophoretic mobility shift assays (EMSAs)

Full-length DIVARICATA1 was cloned into the pMAL-c2x vector and fused in-frame with the Maltose binding protein (MBP) tag. The constructs were transformed into *E. coli* strain BL21, which was then grown in liquid medium to an OD at 600 nm of 0.4, treated with 1 mM IPTG to induce expression, and grown for a further 16 h at 16 °C with shaking at 130 rpm. The recombinant protein was purified with amylose beads according to the manufacturer’s instructions (NEB, USA).

The EMSA was carried out as previously described, with minor modifications (Sun *et al*., 2020). The *CCS* promoter probe DNA containing the wild-type and mutated *cis*-element probes was labelled using a Pierce Biotin 3’ End DNA Labelling Kit (Thermo Fisher Scientific, USA). EMSA was performed using a LightShift Chemiluminescent EMSA Kit (Thermo Fisher Scientific, USA) according to the manufacturer’s instructions.

#### ChIP-PCR

The recombinant DIVARICATA1 protein were used as antigens for the polyclonal antibody production, the antibodies were produced in rabbit as described previously (Zhu *et al*., 2019). ChIP-PCR was performed as described previously, with some modifications (Zhu *et al*., 2019). Briefly, pepper pericarp samples were collected from inbred line 59 fruit at developmental stage 38 DAP. Five-gram samples were crosslinked on ice for 30 min under vacuum in crosslinking buffer. The chromatin was sonicated into 250~500 bp fragments on ice. Subsequently, the solubilized chromatin was immunoprecipitated with the DIVARICATA1 polyclonal antibody or mouse IgG. Finally, the recrosslinked and precipitated DNA was purified, and quantitative real-time PCR analyses were performed to analyse the enrichment. The primers used in this study are listed in Table S2.

### ABA treatment and sampling

The concentration and application of ABA were as described in previous studies with slight modifications (Casey Barickman *et al*., 2014; Cervantes-Hernández *et al*., 2022; Liu *et al*., 2017). Briefly, mature green (30 DPA) pepper fruit of the inbred line ‘59’ was treated with 50 μM ABA (Sigma-Aldrich, USA). The fruits were sampled at the indicated time points (0 h, 0.5 h, 1 h, 3 h, 6 h, 12 h and 24 h) for gene expression analysis, and 38 DPA fruits were sampled for capsanthin analysis. To evaluate *DIVARICATA1* -silenced plants in response to ABA, VIGS control plants and *DIVARICATA1* -silenced mature green (30 DPA) fruits were treated with 50 μM ABA. After ABA treatment for 6 h, fruits were sampled for gene expression analysis, and 38 DPA fruits were sampled for capsanthin determination.

## Acknowledgement

This work was supported by the National Natural Science Foundation of China (32102380, U21A20230, 32070331 and 32072580). Guangdong Basic and Applied Basic Research Foundation (2022A1515012547).

## Author contributions

ZZS and JJL conceived and supervised the project. ZSZ designed the study, performed key experiments, and wrote the manuscript. HH provided some constructive suggestions for the research and revised the manuscript. ZYN, JLS, BMS, JLW and CJC performed some experiments. XJZ revised the manuscript. NW, BHC, MXC, KHC, GJC, YTC and JD provided some useful suggestions regarding this study.

## Supplementary information

Supplementary Figures and Tables

